# Expanding primary cells from mucoepidermoid and other salivary gland neoplasms for genetic and chemosensitivity testing

**DOI:** 10.1101/171652

**Authors:** Ahmad M. Alamri, Xuefeng Liu, Jan K. Blancato, Bassem R. Haddad, Weisheng Wang, Xiaogang Zhong, Sujata Choudhary, Ewa Krawczyk, Bhaskar V. Kallakury, Bruce J. Davidson, Priscilla A. Furth

## Abstract

Restricted availability of cell and animal models is a rate-limiting step for investigation of salivary gland neoplasm pathophysiology and therapeutic response. Conditionally reprogrammed cell (CRC) technology enables establishment of primary epithelial cell cultures from patient material. This study tested a translational workflow for acquisition, expansion and testing of CRC-derived primary cultures of salivary gland neoplasms from patients presenting to an academic surgical practice. Results showed cultured cells were sufficient for epithelial cell-specific transcriptome characterization to detect candidate therapeutic pathways and fusion genes in addition to screening for cancer-risk-associated single nucleotide polymorphisms (SNPs) and driver gene mutations through exome sequencing. Focused study of primary cultures of a low-grade mucoepidermoid carcinoma demonstrated Amphiregulin-Mechanistic Target of Rapamycin-AKT/Protein kinase B (AKT) pathway activation, identified through bioinformatics and subsequently confirmed as present in primary tissue and preserved through different secondary 2D and 3D culture media and xenografts. Candidate therapeutic testing showed that the allosteric AKT inhibitor MK2206 reproducibly inhibited cell survival across different culture formats. In contrast, the cells appeared resistant to the adenosine triphosphate competitive AKT inhibitor GSK690693. Procedures employed here illustrate an approach for reproducibly obtaining material for pathophysiological studies of salivary gland neoplasms, and other less common epithelial cancer types, that can be executed without compromising pathological examination of patient specimens. The approach permits combined genetic and cell-based physiological and therapeutic investigations in addition to more traditional pathologic studies and can be used to build sustainable bio-banks for future inquiries.

## Introduction

Primary salivary gland neoplasms arise from both major (parotid, submandibular, or sublingual gland) and minor salivary glands (1). They are composed of benign and malignant tumors of different histopathological types and exhibit a range of responses to chemotherapy (2). Relatively uncommon, they comprise approximately 5% of head and neck tumors (3) with an incidence of 1.7 cases per 100,000 individuals in the United States (4). Limited case numbers and access to patient samples coupled with scarce availability of authenticated cell lines and animal models translate into our current relatively restricted understanding of salivary gland cancer pathophysiology and chemotherapeutic response (5). Standard treatment is surgical resection followed by postoperative radiation (6). Chemotherapy is reserved for recurrence, metastases, and when surgical resection is not possible (7). In addition to being a site for primary cancers, the parotid gland also hosts non-salivary metastatic disease, particularly cutaneous squamous cell cancers, pathophysiology attributed to the presence of glandular lymph nodes (8). Sialoadenitis is a non-malignant inflammatory condition that can present as a salivary gland neoplasm (9).

Mucoepidermoid carcinoma (MEC) is the most common malignant salivary gland tumor representing 30-40% of all salivary gland malignancies (10). CREB Regulated Transcription Coactivator 1 (*CRTC1*)-Mastermind Like Transcriptional Coactivator (*MAML2*), *CRTC3*-*MAML2* and EWS RNA Binding Protein 1 (*EWS*)-POU Class 5 Homeobox 1 (*POU5F1*) fusion genes are reported to be more frequent in low as compared to high grade MEC (11–13). Presence of *CRTC1*-*MAML2* is associated with overexpression of the EGF family member Amphiregulin (AREG) (14–18), where the presence of both has been correlated with longer disease-free survival (19).

AREG is an Epidermal Growth Factor (EGF) family member acting through the EGF Receptor (EGFR) to activate Phosphatidylinositol-4,5-bisphosphate 3-kinase (PI3K) and AKT/Protein kinase B (AKT) pathways that promote cell survival and proliferation. EGFR is an ErbB family member proposed as a therapeutic biomarker in head and neck cancer (20). Activated AKT pathways are reported in many different types of salivary gland cancer (21,22) and AKT inhibitors with dissimilar mechanisms of action are available (23,24). MK2206 is an oral allosteric highly selective AKT inhibitor that reduces levels of phosphorylated (p-) AKT while GSK690693 is an ATP-competitive AKT inhibitor where exposure results in increased levels of p-AKT but reduced levels of p-Glycogen Synthase Kinase 3 Alpha/Beta (GSK3α/β) and p-Mechanistic Target Of Rapamycin (mTOR) (25,26) MK2206 has documented activity in phase II solid tumor clinical trials (27–29). GSK690693 is a validated reference molecule for its class (25).

Conditionally reprogrammed cell (CRC) technology can generate primary epithelial cell cultures from normal, benign and malignant tissue from different species (30–35). Here we explored its utility for establishing primary cell cultures from different types of human salivary gland neoplasms, examined if sufficient material for next generation sequencing could be obtained from the initial cultures, and tested if the cells could be expanded to evaluate growth and chemosensitivity under different culture conditions. In human normal keratinocytes the Rho kinase inhibitor (ROCK) Y-27632 induces down-regulation of genes involved in keratinization and differentiation (36) while in malignant mouse mammary epithelial cells use of CRC technology favors up-regulation of keratin (*KRT*) differentiation genes when compared to mammary optimized media (34). In mammary cells this is associated with up-regulation of Tumor Protein (TP)p53 family genes (34,37). Therefore, the study compared expression of TP53 and salivary epithelial cell KRT differentiation-linked proteins (38,39) in original tissue and cells cultured under CRC and non-CRC conditions.

RNA sequencing (RNA-seq) is an established tool for unbiased transcriptome characterization and fusion gene screening (40,41). Whole exome sequencing is a means to evaluate DNA for known driver mutations, single nucleotide polymorphisms (SNPs), and insertions and deletions (indels) (42). RNAseq results from CRC cultured cells and original tissue show high concordance when passage number is limited (34). Here we tested if combining CRC technology with next generation sequencing (NGS) could be used to correctly identify candidate therapeutic pathways present in the parenchyma of the original tissue.

## Methods and Materials

### Primary salivary gland epithelial cell culture, xenograft and cytogenetic analysis

Fresh tissue not required for pathological diagnosis were serially collected over approximately one year from consenting patients with salivary gland neoplasms. Informed consent was obtained by the Non-Therapeutic Subject Registry (NTSR) working under Institutional Review Board approval. Tissue obtained from surgically excised specimens (0.5-1cm^3^ specimen, n=8 paired samples; n=3 single specimens) or fine needle aspirates (n=1 paired sample) was provided to the Histopathology and Tissue Shared Resource (HTSR) where identifiers were removed and unique codes applied before transfer to the Tissue Culture Shared Resource (TCSR) for primary culture. Two specimens from different areas of the pathological samples were provided when sufficient tissue was available and each processed as individual specimens for comparison (n=9 paired samples). Deidentified pathology reports were provided by HTSR for all samples. Primary cells were isolated using 1x collagenase/hyaluronidase solution (ThermoFisher, Waltham, MA) followed by 0.25% trypsin-EDTA (ThermoFisher) from fresh tissue (surgical specimens) or placed directly into culture without processing (fine needle aspirates) in complete F medium [Ham’s F-12 nutrient mix (25%) (ThermoFisher) supplemented with 25ng/mL hydrocortisone, 5μg/mL insulin, 0.1nmol/L cholera toxin (Sigma-Aldrich, St Louis, MO), 0.125ng/mL epidermal growth factor, 10μg/mL gentamicin (ThermoFisher), 250ng/mL Fungizone (ThermoFisher), 5μmol/L ROCK inhibitor Y-27632 (Y) (Enzo Life Sciences, Farmingdale, NY)] and 74% complete Dulbecco’s modified Eagle’s medium (DMEM) [10% fetal bovine serum (FBS), 100μg/mL penicillin, 100μg/mL streptomycin, 100μg/mL glutamine (ThermoFisher)] in the presence of irradiated Swiss 3T3-J2 mouse fibroblast feeder cells at 37°C with 5% CO_2_ (30,31,43) and assigned a unique Georgetown University Medical Center (GUMC) primary cell culture identifier. For secondary passage, primary cells were separated from feeder layers by differential trypsin treatment (30s, 0.05% trypsin, ThermoFisher) followed by wash with 1X Phosphate Buffered Saline (PBS) to remove detached feeder layer, second treatment with 0.05% trypsin to detach epithelial cells before placement in conditioned medium + 5μmol/L Y (CM+Y), conditioned medium without Y-27632 (CM), EpiC [EpiCult™-C Human Medium Kit containing 5mL EpiCult™-C Proliferation Supplement (Human), hydrocortisone (10^−6^M)] (Stemcell Technologies, Vancouver, BC), 100μg/mL glutamine and 100μg/mL streptomycin/penicillin (ThermoFisher), or MammoCult™ Human Medium Kit supplemented with MammoCult™ Proliferation Supplement (Human) 50mL, hydrocortisone (10^−6^M), 2μg/mL Heparin solution (Stemcell Technologies, Vancouver, BC), and 100μg/mL streptomycin/penicillin (ThermoFisher). Conditioned medium was prepared by plating irradiated Swiss 3T3-J2 mouse fibroblast feeder cells (7 X 10^6^/T175cm^2^ tissue culture flask) in complete F medium with collection of supernatant F medium (CM) followed by filtration through 0.22-mm pore-size filter (EMD Millipore, Billerica, MA) after three days with storage at −80°c and before use mixed with complete F medium (75%CM/25% complete F) (44). Primary cell cultures were viably frozen and re-cultured when needed. For 3D, embedded cultures in Matrigel (BD Biosciences, Franklin Lakes, NJ), cells were grown to 80-90% confluency, detached by trypsinization, resuspended in complete F medium + Y (4×10^4^ cells/suspension) on ice, mixed with 1.2ml of Matrigel, plated on the surface of precoated 6 well culture plates with two ml of F medium + Y added on top and placed at 37°C with 5% CO_2_ (45). To precoat plates, prechilled 6 well culture plates were coated with a thin layer of Matrigel (200μl) and incubated at 5 min at 37°C. Cultures were maintained for 10 days. For 3D spheroid culture, 1.5 ×10^4^ cell /100μl of CM+Y, CM, and EpiC was plated in 96 well round bottom low attachment plates (Corning, NY, USA), centrifuged at 1000 rpm (5 min), and placed at 37°C with 5% CO_2_. Vybrant™ Cell-Labeling Solution (ThermoFisher) was used to label cells with green fluorescence (10μl cell-labeling solution/1mL cell suspension with incubation at 37°C (30 min) followed by centrifugation at 150rpm (5 min), removal of supernatant followed by washing three times in warm media and resuspension. Digital images of 2D and 3D primary cell cultures were taken using EVOS™ XL Core Cell Imaging System (ThermoFisher). For xenograft development, six-week-old athymic nude (*NU (NCr)*-*Foxn1nu*) female mice (n=20) (Harlan Laboratories, Inc., Frederick, MD) (26-30gm) were housed in barrier zones in single-sex sterilized ventilated cages at Georgetown University and acclimatized one week prior to primary cell inoculation (10^6^ /site in 25μl PBS/25μl Matrigel (BD Biosciences, Franklin Lakes, NJ) via 1cc syringe/27-gauge needle) into thoracic and inguinal mammary fatpads (4 sites/mouse) and into flank subcutaneously (2 sites/mouse) through a skin incision under isoflurane anesthesia. Mice were monitored weekly with measurement of palpable and visible tumors and euthanized by CO_2_ inhalation followed by cervical dislocation when mice reached 6 months of age or palpable tumors > 1cm^3^, whichever occurred first. At necropsy mice were examined for xenograft growth at injected sites and tissue removed for formalin fixation and processing for pathological examination. All animal procedures were performed and approved by the GUMC Institutional Animal Care and Use Committee. For conventional cytogenetic analysis (karyotyping), primary GUMC220/221 cells were cultured in F media + Y (passage (P) 9 for both). Chromosome preparation and G-banding assays were performed using standard protocols (46) and chromosomes identified and classified according to standard cytogenetic nomenclature (47).

### RNAseq, Exome Sequencing and Analyses

Primary cells cultured in CRC were pelleted and total RNA (n=14) (RNeasy Mini Kits, Qiagen, Valencia, CA) and DNA (n=8) (MasterPure complete DNA and RNA purification Kit, Epicentre, Madison, WI) extracted and quality analyzed (Nanodrop, Agilent Bioanalyzer 2100, Agilent Technologies, Santa Clara, CA). For RNAseq shotgun library construction (200bp insert) was completed and then sequenced using a HiSeq2000 (91bp pair-ended lane generating 2 Gb/sample, raw data) (Macrogen, Seoul, South Korea) and aligned to a human reference genome (UCSChg19). Transcriptomes were assembled, transcript abundance estimated (FPKM: fragments per kilobase of transcript per million mapped reads) and statistically significantly differentially expressed genes (DEGs) between paired specimens determined (2fold up- or 0.5 down-regulated expression) (Cufflinks and Cuffdiff) (48,49). Top ten Hallmark gene sets were identified using (Molecular Signatures Database (MSigDB) and Ingenuity Pathway Analysis (IPA) for top ten canonical pathways for DEGs ≥500 FPKM. Heat maps were generated using ClusVis (a web tool for visualizing clustering data (BETA) (50). Candidate fusion genes were identified with FusionCatcher Software (fusioncatcher.py 0.99.3e beta) (41). For Human Exome Capture sequencing, the Sure Select Target Enrichment System Capture Process was used with Illumina HiSeq2000 (91bp paired end sequencing with SureSelect Human All Exon V4 (51M) kit (50X on target coverage) followed by analysis of exome sequencing data using Burrows-Wheeler Aligner (51) to map data against UCSChg19 and SAMTOOLS (52) for identification of SNPs and indels. RNAseq files to be uploaded to Gene Expression Omnibus (GEO) and DNA exome sequencing files to Sequence Read Archive (SRA).

### Reverse Transcriptase Polymerase Chain Reaction (RT-PCR), Quantitative RT-PCR and Fluorescence in situ hybridization (FISH)

RNA was extracted from cell pellets (RNeasy Mini Kit, QIAGEN, Gaithersburg, MD), quantified (Nanodrop, Thermo Fisher Scientific) and 1μg total RNA used to prepare cDNA (iScript™ cDNA Synthesis Kit, Bio-Rad). RT-PCR performed using specific primers sets (Supplementary Tables 2, 3) and GoTaq^®^ Green Master Mix (Promega, Madison, WI). (30-40 cycles: 1min: 95°C, 45 S-1min: 55-60°C, 2min: 72°C) and products visualized using ethidium bromide staining after electrophoresis on agarose gels (Bio-Rad Universal Hood II, Bio-Rad, Hercules, CA). Candidate PCR products were extracted (QIAquick^®^ Gel Extract Kit, QIAGEN, Gaithersburg, MD) and sequenced (GENEWIZ, South Plainfield, NJ). For qRTPCR, 1μg total RNA was converted to cDNA using Taqman reverse transcription reagents (Applied Biosystems, ThermoFisher) and 2.5μl of the resulting cDNA used for each 20μl reaction volume containing 10μl of 2x TaqMan^®^ Fast Universal PCR Master Mix (Applied Biosystems), 6.5μl of nuclease-free water, and 1μl of Taqman primers for *AREG*, (4331182, ThermoFisher) or control 18s (ThermoFisher). Applied Biosystems StepOneTM/ StepOnePlusTM Real-Time PCR System (Applied Biosystem) was used with 2min: 50°C, 10min: 95°C, and 40 cycle (15s: 95°C, 1min:60°C). Data analyzed by StepOne™ Software 2.3 (Applied Biosystems). For FISH, GUMC220/221 primary cells cultured in F medium+Y-27632 were detached from flasks by trypsinization with 0.05% trypsin and treated with hypotonic solution (0.075M KCl), fixed on slides (3:1 methyl alcohol and glacial acetic acid), pretreated (Nonidet P-40, 20x saline-sodium citrate (SSC), and distilled water) at 37°C for 30 and then dehydrated in an ascending series of ethanol solutions (70%, 80%, 95%) for 2 min each. FFPE sections were baked on slides overnight at 60 °C to adhere tissue, deparaffinized by consecutive 10-min xylene washes before being rehydrated through 100% ethanol incubation for 10 min at RT. Tissues on the slides were permeabilized and digested using the Abbott tissue digestion kit containing pepsin (Naperville, IL), according to manufacturer’s instruction, and washed. Probes for *KRT14*-*20*-*RE* (Empire Genomics, Buffalo, NY) covering 180 Kb of chromosome 17q21.2 including the entire *KRT14* gene, and *KRT5*-*20*-GR (Empire Genomics, Buffalo, NY) covering 151Kb of chromosome12q13.13 including the entire *KRT5* gene, were denatured at 74 ° C for 10 minutes, mixed with hybrdization buffer and immediately transferred to an ice bucket. Denatured probes were added to sections, slides sealed with rubber cement and co-denatured for 8 min on a HYBrite heat plate (Vysis, Naperville, Illinois) at 85 ° C for 8 min, and then incubated for 16 to 24 hrs at 37 ° C. Coverslips were then removed and slides were washed in 2X SSC hybridization buffer for 2 min at 73°C and transferred to 2X SSC at RT for 5 min. Slides were air dried in the dark for 1 hr in an upright position and counterstained with 4’, 6-diamidino-2-phenylindole (DAPI) (Vector Laboratories, Inc. Burlingame, CA) and viewed on an Axioscope fluorescence microscope (Zeiss, Oberkochen, Germany) and imaged with Applied Imaging Cytovision software (Pittsburgh, PA).

### Histology, immunohistochemistry and western blotting

For hematoxylin and eosin (H&E) and immunohistochemistry (IHC) examination, FFPE five μm sections of excised original tissue (T) with identifiers removed, xenografts (X) and Matrigel cultures (M) and primary cell pellets (P) prepared after fixation in 10% buffered formalin (Fisher Scientific, Hampton, NH) overnight at 4°C were obtained from HTSR. For IHC, antigen retrieval was performed using 10 mM Citrate pH 6.0 buffer, EDTA pH 8.0 (home made), Tris/EDTA pH 9.0 (Genemed, South San Francisco, CA), and /or Envision FLEX Target Retrieval Solution Low pH (Agilent, Santa Clara, CA). For western blotting (WB), cell lysates were collected using radioimmunoprecipitation assay buffer (RIPA) (ThermoFisher) supplemented with Halt Protease and Phosphatase Inhibitor Cocktail (1:100) (ThermoFisher), sonicated, clarified by centrifugation (10min), concentration measured (Pierce^®^ 660 nm Protein Assay Reagent, ThermoFisher) and 8 μg protein loaded with 2X Laemmli Sample Buffer (Bio-Rad, Hercules, CA) with 0.05 2-Mercaptoethanol (BME) on 10% gels (Bio-Rad) followed by transfer to PVDF membrane (Bio-Rad, Hercules, CA). Membranes were blocked with 7% bovine serum albumin (BSA) (Santa Cruz Biotechnology) prepared in 1xTris-buffered saline, TWEEN 20 (TBST) (100mL of 10x TBS (Bio-Rad) + 1ml of 100% Tween 20 (Fisher) + 890 distilled water) for 1 hr RT, incubated overnight at 4°C with primary antibodies in 5% BSA in 1xTBST, washed three time with 1xTBST, followed by incubation with appropriate polyclonal horseradish peroxidase-conjugated (HPRC) secondary antibodies (1:5000 dilution in 1xTBST) for 1 h at room temperature, visualized by chemiluminescence (HyGLO Chemiluminescent HRP Antibody Detection Reagent, Denville Scientific, South Plainfield, NJ). Primary antibodies: Amphiregulin (AREG) (HPA008720, Sigma-Aldrich, St. Louis, MO, dilutions: T, X-1:150, P-1:600, M-1:75). Epidermal Growth Factor Receptor (EGFR) (ab52894, Abcam, Cambridge, MA, dilutions T-1:75, P-1:200, X-1:200, M-1:100). p-EGFR (2236, Cell Signaling, dilutions T-1:00, P-1:1000, X-1:150, M-1:100). Pan-AKT (4691, Cell Signaling, dilution WB-1:1000). p-AKT (Ser473) (4060, Cell Signaling, dilutions T,P,X,M-1:30, WB-1:2000. p-Akt (The308) (13038, Cell Signaling, dilutions WB-1:1000). mTOR (2972, Cell Signaling, dilutions WB: 1:1000). p-mTOR (2976, Cell Signaling, dilutions T,P,X,M-1:150). p-mTOR (5536, Cell Signaling, dilution WB-1:1000). GSK3α/β (5776, Cell Signaling, dilution WB-1:1000). p-GSK3α/β, 9331, Cell Signaling, dilution WB-1:1000). β-Actin (3700, Cell Signaling: WB-1:1000). Cytokeratin 5 (KRT5) (PRB-160P, BioLegend, San Diego, CA, dilutions T-1:150, P-1:2000, M-1200). Cytokeratin 14 (KRT14) (PRB-155P, BioLegend, dilutions: T-1:150, P-1:2000, M-100). Cytokeratin 8 (KRT8) (ab59400, Abcam, dilutions: T,P-1:75, M-1:100). Cytokeratin 18 (KRT18) (4548, Cell Signaling: T,P-1:200, M-1:200). p53 (OP33, Calbiochem-EMD Millipore, Billerica, MA, dilutions: T-1:40, P,M-1:40). Human mitochondrial protein (clone 113-1, EMD Millipore, dilutions: X-1:100). Secondary antibodies: WB, HRP antibodies 1:5000 (Santa Cruz Biotechnology); IHC, ABC or HRP antibodies. A board certified pathologist (BVK) read histology and IHC blinded to antibody identity. Histology images taken using a Nikon Eclipse E800 Microscope/ NIS-Elements BR 4.30.02 64-bit software (Nikon Instruments Inc., Melville, USA). Western blot images taken by an Amersham™ Imager 600 (GE Health Care Life Sciences, Pittsburg, PA) and quantified using ImageJ (http://imagei.nih.gov.proxy.library.georgetown.edu/ii). Relative expression levels were determined after normalizing total protein to actin signals obtained from the same membrane and p-protein to corresponding total protein. Means and Standard errors of the mean were calculated and data statistically analyzed and plotted using Prism 6.0 (GraphPad Software, La Jolla, CA). Each western blot was performed three times.

### Chemosensitivity and cell survival analyses

GUMC220/221 and MDA-MB-453 cells were plated in 96 well plates (5 ×10^4^ cell /100μl of CM+Y, CM, or EpiC) for either 2D (VWR, Radnor, PA) or 3D (round bottom low attachment plates) culture. For 2D culture, cells were incubated overnight at 37°C with 5% CO_2_ followed by removal of initial plating media and replacement with media containing different concentrations of MK2206 2HCI (MK2206) (1.2μM-20μM), GSK690693 (2.5μM-40μM), doxorubicin (0.3μM-5μM) or vehicle-only Dimethyl sulfoxide (DMSO) control. For 3D culture, cells were incubated overnight at 37°C with 5% CO_2_ followed by addition of MK2206 (1.2μM-40μM), GSK690693 (1.2μM-40μM), doxorubicin (0.3μM-10μM) or vehicle-only (DMSO) control to the media with final concentration calculated for total well volume. Stock solutions (10mM) of MK2206, GSK690693, and doxorubicin (Selleckchem, Huston, TX), was prepared in DMSO (VWR, Radnor, PA). Cell viability was measured using CellTiter-Glo^®^ Luminescent Cell Viability Assay (Promega, Madison, WI) occurring to the manufacturer’s instruction after three (2D cultures) or five (3D cultures) days using Veritas microplate luminometer turner biosystems and GloMax^®^-96 Microplate Luminometer Software (Promega). Each experiment included three cell/condition technical replicates and each experiment was completed three times. DMSO luminescence values were normalized to non-treated wells and drug-treated wells normalized to corresponding DMSO wells.

### Statistics

Fishers exact test was used to compare frequency distribution of top ten HALLMARK pathways, unpaired t test, two-tailed to compare AREG qRTPCR, One-way ANOVA test was used to compare drugs response treatment on primary cells, unpaired t-test, Welch-corrected, one-tailed was used for western blot analysis and means and standard errors of the mean (SEM) calculated and plotted (GraphPad Software Prism 6.0, Inc. La Jolla, CA).

## Results

### Primary cultures from salivary gland neoplasms established using CRC technology were expanded under alternate culture conditions and yielded sufficient material at low passage numbers for NGS analyses

Primary cultures from malignant (MEC, carcinoma ex pleomorphic adenoma (ca ex PA), squamous cell carcinoma (SCC), squamous cell carcinoma metastatic to salivary gland (metastatic SCC), diffuse large B cell lymphoma in salivary gland and benign (pleomorphic adenoma (PA), benign ductal squamous metaplasia) salivary gland neoplasms were established using CRC technology (Table 1). Specimens were obtained from surgically excised tissue with the exception of ca ex PA (attained from fine needle aspirates). Paired independently derived cultures were established from different geographic regions of the same tumors (n=9) to be used for NGS and secondary culture studies (Figure 1A). Formalin fixed paraffin embedded (FFPE) parent tissue was available for validation studies. Neither age nor sex influenced likelihood of culture establishment however no cultures could be established from the two sialoadenitis cases attempted. The well differentiated MEC (GUMC220/221) and two metastatic to salivary gland SCC specimens (GUMC264/265 and GUMC367) were selected for secondary culture studies (Table 2). All cells expanded for multiple passages (3–15) under the three secondary 2D culture conditions attempted (Figure 1B) including the well-differentiated low-grade MEC (Figure 1C). Paired cultures demonstrated similar secondary culture kinetics but expansion was fastest using conditioned media (CM) + Y-27632 (Y) (Figure 1D). Karyotype was normal for the GUMC220/221 cells grown in CRC (Figure 1E). While all five cultures propagated in alternative media including EpiCult™-C (EpiC) (StemCell Technologies, Vancouver CN), only GUMC220/221 formed xenografts (Table 2, Figure 2A). GUMC220/221 also propagated in MammoCult™ (StemCell Technologies) and formed spheroids when grown in Matrigel and on low adherent plates with both CM+Y and EpiC media (Figure 2B,C). Sufficient cells for NGS were obtained after two (GUMC220/221, GUMC264/265, GUMC299, GUMC367, GUMC332/349, GUMC311, GUMC436/446, GUMC572/573), three (GUMC374/378, GUMC312) and four (GUMC284) passages. The magnitudes of statistically significant differentially expressed genes (DEGs) were less than 0.4% of genes expressed for all seven paired cultures examined (range 0-0.38%) (Table 3). Hierarchical clustering demonstrated limited dissimilarity between paired specimens (Supplementary figure 1) but all cultures demonstrated significant expression of *KRT* genes consistent with epithelial origin. No known pathogenic cancer mutations or driver genes were identified; however, cancer risk-related probable pathogenic SNPs were reproducibly present in both of the four paired samples subjected to exome sequencing (Table 4). In GUMC220/221 these included *Brca2 rs144848*, *TP53* rs1042522, *AURKA* rs2273535, *RET* rs1800858 and *ADH1B* rs1229984.

**Figure 1.**
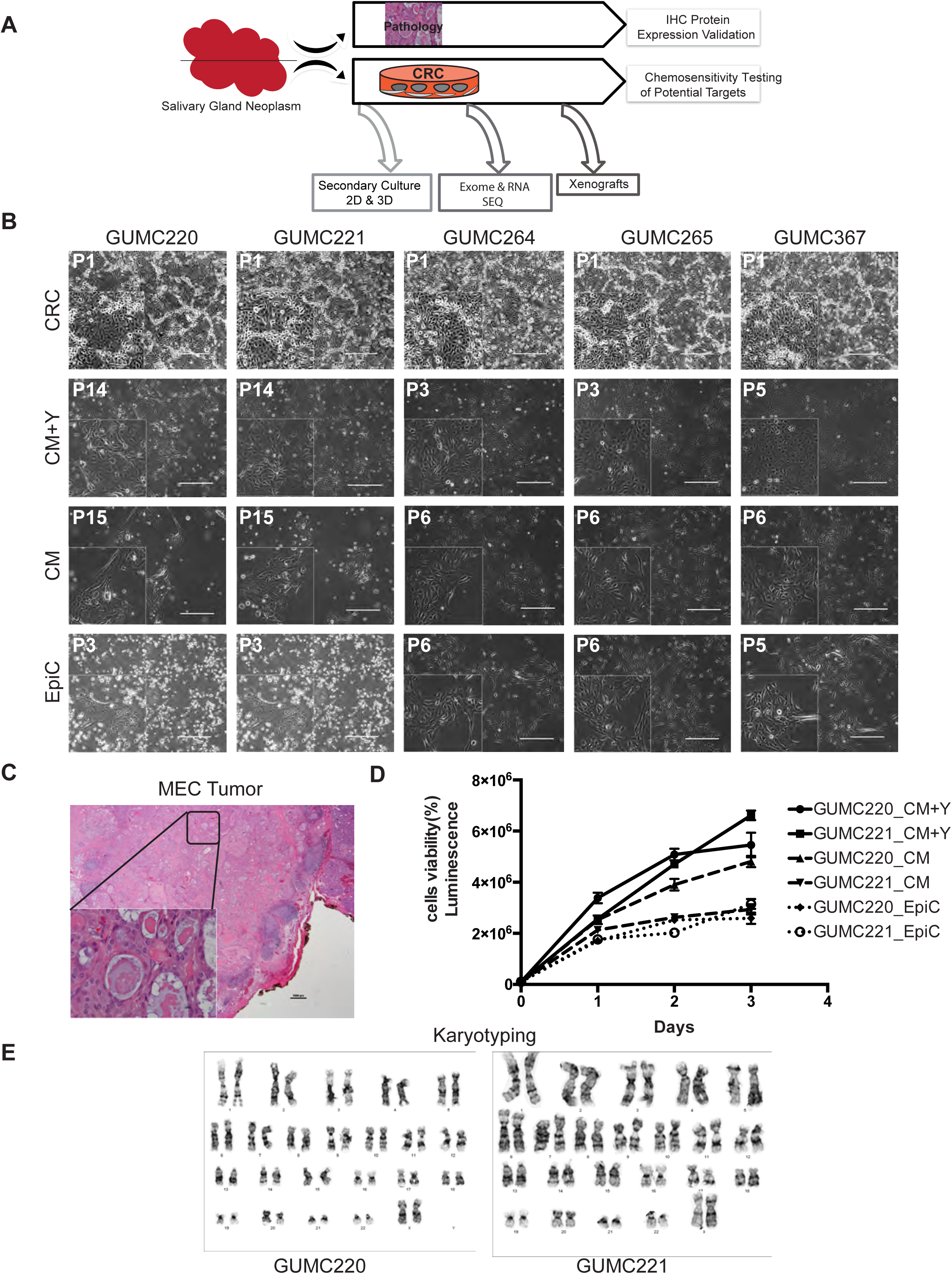
Primary cancer cell cultures established in CRC were secondarily propagated without the presence of the rho kinase inhibitor or conditioned media. (A) Overall experimental design. (B) Representative phase-contrast images of primary GUMC220/221 derived from a well-differentiated sublingual mucoepidermoid carcinoma (MEC) in CRC, CM+Y, CM, and EpiC media. GUMC264/265 and GUMC367 cultures derived from squamous cancers metastatic to parotid gland also grew in all three medias. Passage number indicated top right. Images taken at 10x. Size bars = 400μm. (C) H&E image of the MEC tumor from which GUMC220 and GUMC221 cultures were derived. Image taken at 1x. Size bar = 10μm. Bottom left inset shows magnified image taken at 40x. Size bar = 10μm. (D) Comparison of GUMC220 and GUMC221 cell viability curves in CM+Y, CM, and EpiC media over three days. Luminescence: CellTiter-Glo^®^, Promega, Madison, WI, USA. (E) Primary CRC-cultured GUMC220 and GUMC221 cells show normal karyotypes (46, XX). CRC, conditionally reprogramming cells; CM+Y, conditioned media + ROCK inhibitor (Y-27632); CM, conditioned media without Y-27632; EpiC, EpiCult™-C Human Medium.

**Figure 2.**
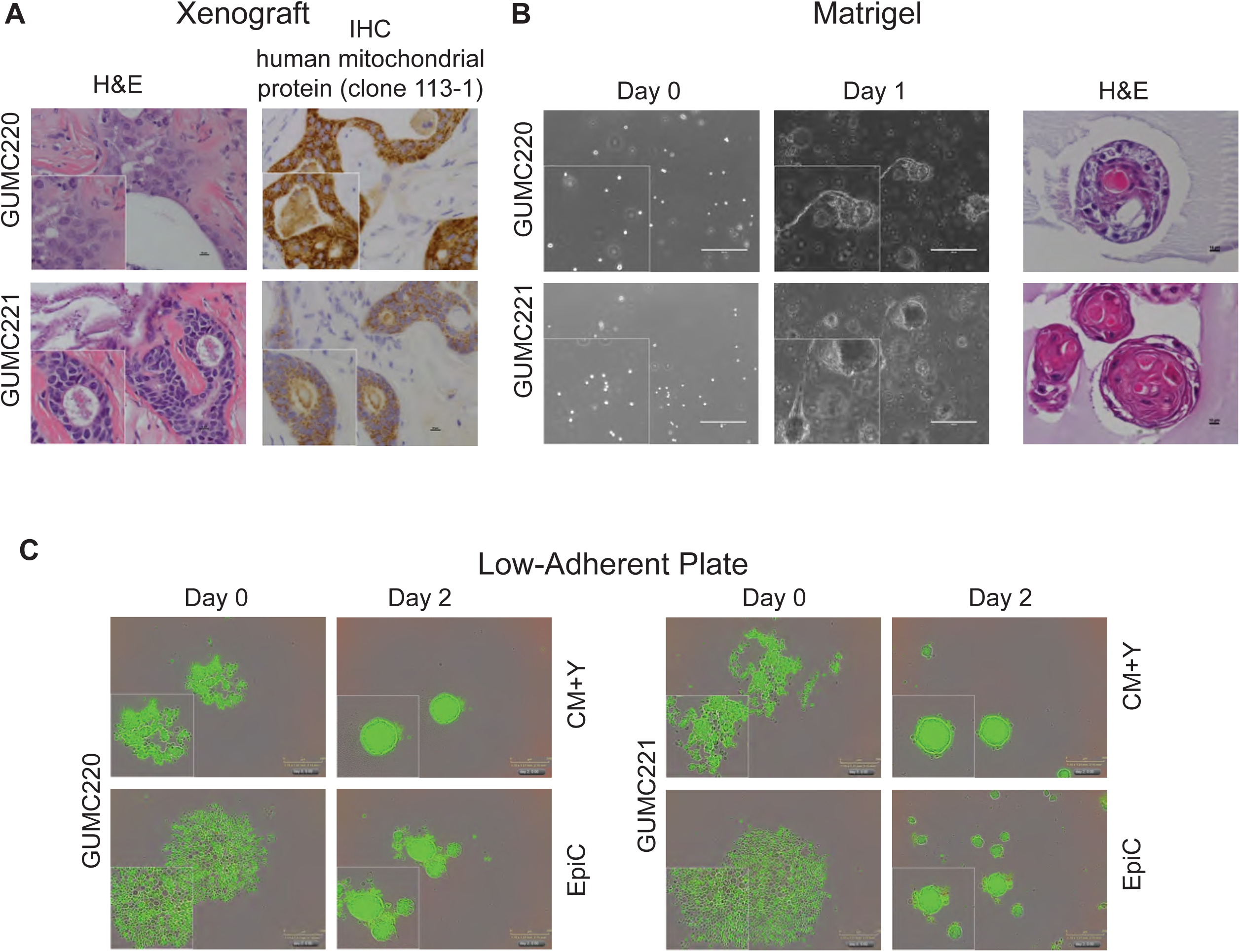
GUMC220 and GUMC221 mucoepidermoid carcinoma primary cells demonstrated xenograft growth *in vivo* and 3D growth *in vitro*. (A) Representative images of xenograft histology: H& E (left), IHC for human mitochondrial protein (clone 113-1) (right). Images taken at 40x. Size bar = 10μm. Bottom left inset shows magnified images. (B) Representative phase contrast images from *in vitro* 3D Matrigel culture: single cell suspensions at day 0 (left), spheroid formation at day 7 (middle), and representative H&E images of spheres (right). Images taken at 10x. Size bar = 400μm. Bottom left inset shows magnified images. (C) Representative green fluorescence images from *in vitro* 3D low-adherent plate culture: single cell suspensions in CM+Y and EpiC at day 0 (left), spheroid formation in CM+Y and EpiC at day 2 (right). Images taken at 10x. Size bar = 300μm. Bottom left insert shows magnified images.

**Table 1:** Primary cell cultures established from salivary gland tissue.

| **Pathology** | **Salivary Gland** | **Race** | **Age** | **Sex** | **Primary Culture Identifiers** |
| --- | --- | --- | --- | --- | --- |
| **Mucoepidermoid carcinoma** | Sublingual gland | White/Caucasian | 56 | F | GUMC220<br>GUMC221 |
| **Carcinoma ex pleomorphic adenoma** | Parotid gland | Unknown | 55 | M | GUMC374<br>GUMC378 |
| **Squamous cell carcinoma** | Sublingual gland | White/Caucasian | 65 | F | GUMC366<br>GUMC360 |
| **Metastatic poorly differentiated squamous cell carcinoma** | Parotid gland | White/Caucasian | 70 | M | GUMC264<br>GUMC265 |
| **Metastatic squamous cell carcinoma** | Parotid gland | White/Caucasian | 80 | M | GUMC367 |
| **Benign pleomorphic adenoma** | Parotid gland | White/Caucasian | 36 | F | GUMC572<br>GUMC573 |
| **Benign pleomorphic adenoma** | Parotid gland | White/Caucasian | 39 | M | GUMC349<br>GUMC332 |
| **Benign pleomorphic adenoma** | Parotid gland | Other | 39 | M | GUMC436<br>GUMC446 |
| **Benign pleomorphic adenoma** | Parotid gland | Unknown | 45 | F | GUMC520 |
| **Benign pleomorphic adenoma** | Submandibular gland | White/Caucasian | 68 | M | GUMC299<br>GUMC284 |
| **Diffuse large B cell lymphoma** | Submandibular gland | Black/African American | 74 | F | GUMC311<br>GUMC312 |
| **Benign ductal squamous metaplasia** | Submandibular gland | Unknown | 31 | M | GUMC224 |

M: Male. F: Female. GUMC: Georgetown University Medical Center.

**Table 2:**
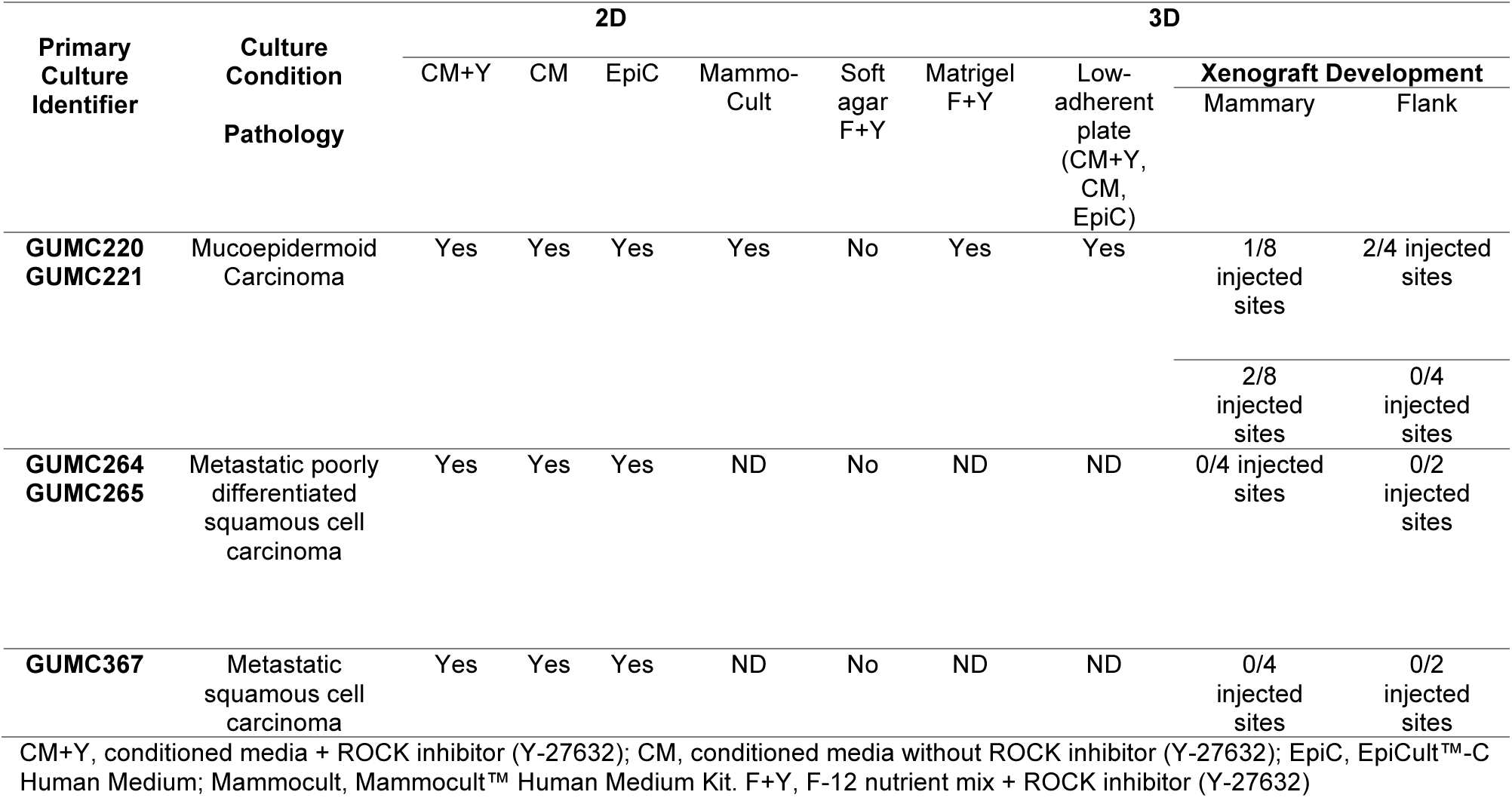
Secondary *in vitro* culture and xenograft outcomes.

**Table 3:** Number of statistically significantly differentially expressed genes (DEGs) between paired primary culture samples from two different geographical regions of tumors.

| Primary Culture Identifier | Number of DEGs Significantly Upregulated<sup>1</sup> | Total Number Significant DEGs | Total number of genes >5 FPKM | Pathology |
| --- | --- | --- | --- | --- |
| **GUMC220** | 48 | 66 | 26006 | Mucoepidermoid carcinoma |
| **GUMC221** | 18 |  |  |  |
| **GUMC264** | 0 | 0 | 26006 | Metastatic poorly differentiated squamous cell carcinoma |
| **GUMC265** | 0 |  |  |  |
| **GUMC367** | 0 | 0 | 26006 | Metastatic squamous cell carcinoma |
| **GUMC374** | 0 | 0 | 26006 | Carcinoma ex pleomorphic adenoma |
| **GUMC378** | 0 |  |  |  |
| **GUMC572** | 19 | 84 | 26006 | Benign pleomorphic adenoma |
| **GUMC573** | 65 |  |  |  |
| **GUMC332** | 0 | 18 | 26006 | Benign pleomorphic adenoma |
| **GUMC349** | 18 |  |  |  |
| **GUMC436** | 57 | 98 | 26006 | Benign pleomorphic adenoma |
| **GUMC446** | 41 |  |  |  |
| **GUMC284** | 28 | 68 | 26006 | Benign pleomorphic adenoma |
| **GUMC299** | 40 |  |  |  |
| **GUMC311** | 12 | 14 | 26006 | Diffuse large B cell lymphoma |
| **GUMC312** | 2 |  |  |  |

FPKM: Fragments Per Kilobase of transcript per Million mapped reads. GUMC: Georgetown University Medical Center

**Table 4:**
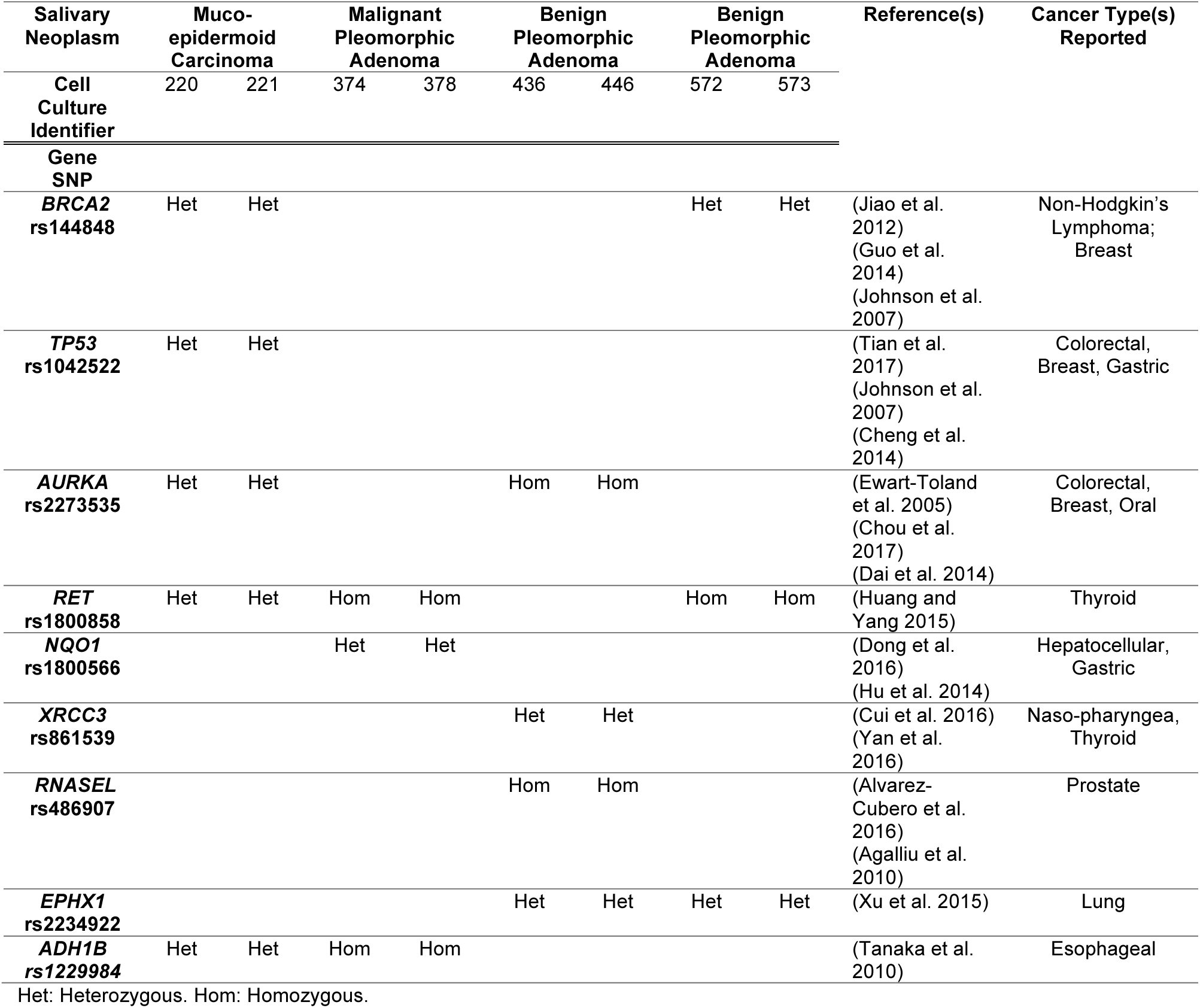
Single nucleotide polymorphisms (SNPS) with published association with increased cancer risk

| Salivary Neoplasm | Muco-epidermoid Carcinoma |  | Malignant Pleomorphic Adenoma |  | Benign Pleomorphic Adenoma |  | Benign Pleomorphic Adenoma |  | Reference(s) | Cancer Type(s) Reported |
| --- | --- | --- | --- | --- | --- | --- | --- | --- | --- | --- |
| Cell Culture Identifier | 220 | 221 | 374 | 378 | 436 | 446 | 572 | 573 |  |  |
| Gene SNP |  |  |  |  |  |  |  |  |  |  |
| **BRCA2**<br>**rs144848** | Het | Het |  |  |  |  | Het | Het | (Jiao et al. 2012)<br>(Guo et al. 2014)<br>(Johnson et al. 2007) | Non-Hodgkin's Lymphoma;<br>Breast |
| **TP53**<br>**rs1042522** | Het | Het |  |  |  |  |  |  | (Tian et al. 2017)<br>(Johnson et al. 2007)<br>(Cheng et al. 2014) | Colorectal, Breast, Gastric |
| **AURKA**<br>**rs2273535** | Het | Het |  |  | Hom | Hom |  |  | (Ewart-Toland et al. 2005)<br>(Chou et al. 2017)<br>(Dai et al. 2014) | Colorectal, Breast, Oral |
| **RET**<br>**rs1800858** | Het | Het | Hom | Hom |  |  | Hom | Hom | (Huang and Yang 2015) | Thyroid |
| **NQO1**<br>**rs1800566** |  |  | Het | Het |  |  |  |  | (Dong et al. 2016)<br>(Hu et al. 2014) | Hepatocellular, Gastric |
| **XRCC3**<br>**rs861539** |  |  |  |  | Het | Het |  |  | (Cui et al. 2016)<br>(Yan et al. 2016) | Naso-pharyngea, Thyroid |
| **RNASEL**<br>**rs486907** |  |  |  |  | Hom | Hom |  |  | (Alvarez-Cubero et al. 2016)<br>(Agalliu et al. 2010) | Prostate |
| **EPHX1**<br>**rs2234922** |  |  |  |  | Het | Het | Het | Het | (Xu et al. 2015) | Lung |
| **ADH1B**<br>**rs1229984** | Het | Het | Hom | Hom |  |  |  |  | (Tanaka et al. 2010) | Esophageal |

Het: Heterozygous. Hom: Homozygous.

### An activated *AREG*-*EGFR*-*AKT* pathway but no *CRTC1*-*MAML2* fusion gene was identified in the well-differentiated mucoepidermoid carcinoma by transcriptome analyses

To identify cancer associated signaling pathways amenable to *in vitro* therapeutic testing, the Molecular Signatures Database (MSigDB) v6.0 (http://software.broadinstitute.org/gsea/msigdb) was interrogated to identify the top ten Hallmark gene sets with significant overlaps with genes expressed ≥500 Fragments Per Kilobase of transcript per Million mapped reads (FPKM) for each paired cultures. The PI3_AKT_Mtor-Signaling Hallmark gene set was uniquely associated with MEC cultures GUMC 220/221 (p<0.05, Fishers exact, Figure 3A). In addition, the top three canonical pathways identified from Ingenuity Pathway Analysis (IPA) of GUMC220/221 (genes ≥500 FPKM) shared AKT as a central regulator (Eukaryotic Translation Initiation Factor (EIF) Signaling, Regulation of eIF-4 and Ribosomal Protein S6 Kinase (p70Sk) Signaling, mTOR Signaling) (Figure 3B).

**Figure 3.**
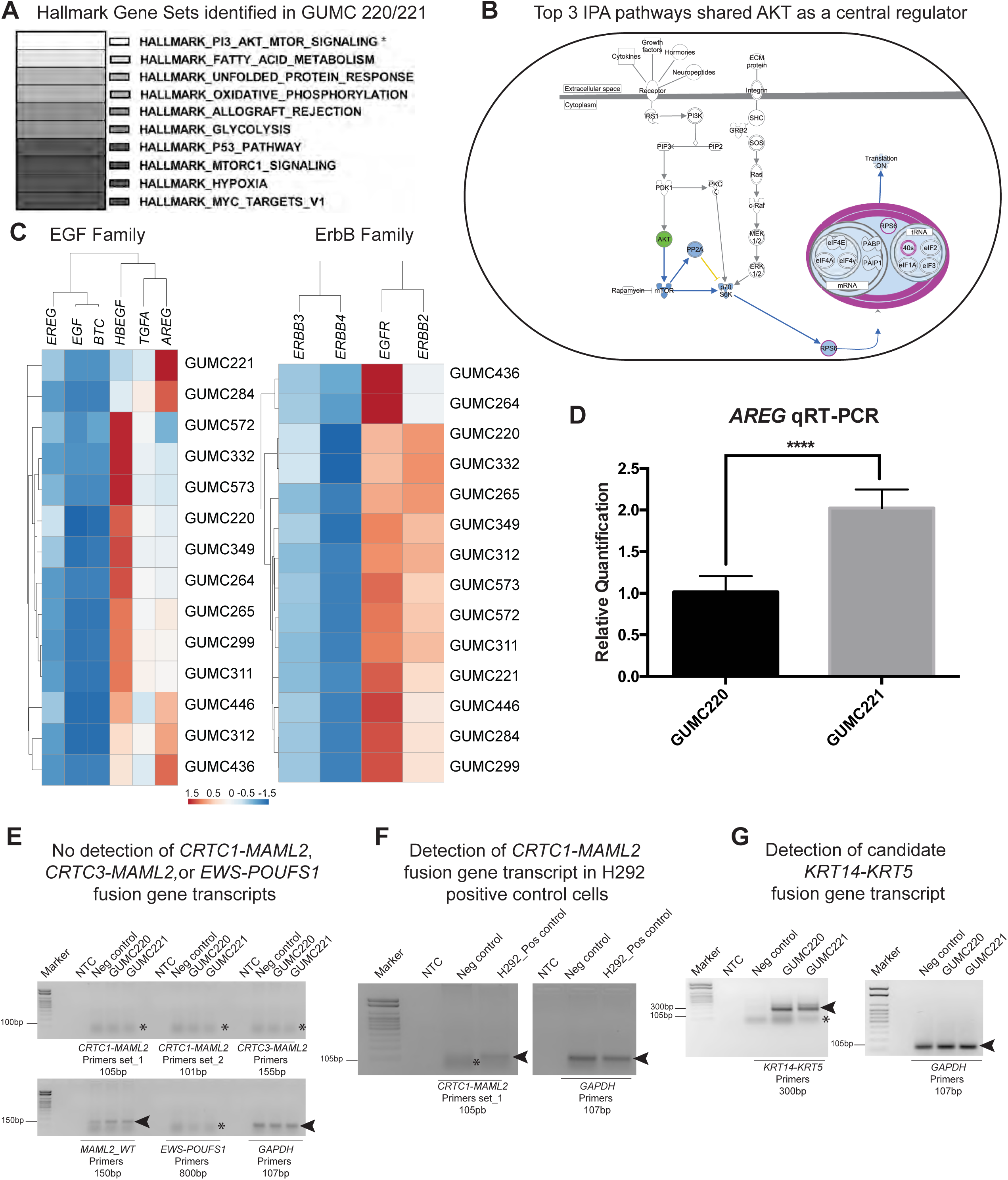
Transcriptome characterization of GUMC220 and GUMC221 mucoepidermoid carcinoma primary cells demonstrated evidence of PI3_AKT_MTOR signaling activation and presence of a candidate KRT14-KRT5 fusion. (A) Top ten Hallmark Gene Sets (MSigDB) identified from genes showing ≥500 FPKM (Fragments Per Kilobase of transcript per Million mapped). Top ten pathways were identical in GUMC220 and GUMC221. Only PI3_AKT_MTOR_Signaling was significantly associated with GUMC220/221 compared to the six other paired culture transcriptomes analyzed (four pleomorphic adenomas: GUMC284/299,332/349,436/446,572/573, one metastatic squamous to parotid: GUMC264/265, one submandibular lymphoma: GUMC311/312, ^*^p<0.05, Fishers exact. (B) Top three canonical pathways identified by Ingenuity Pathway Analyses (IPA) shared AKT as a central regulator. (C) Dendrograms and heat maps of unsupervised hierarchical of FPKM values of genes from EGF (*EREG*, *EGF*, *BTC*, *HBEGF*, *TGFA*, *AREG*) and ErbB (*ERBB3*, *ERBB4*, *EGFR*, *ERRB2*) families. (D) Bar graphs illustrating relative qRT-PCR quantification of *AREG* expression in GUMC220 and GUMC221 cells. Mean±SEM shown, ^****^p<0.0001, t-test, n=3. (E) Ethidium bromide-stained agarose gels showing absence of detection of *CRTC1*-*MAML2*, *CRTC3*-*MAML2*, and *EWS*-POUFS1 fusions (asterisks indicate primer dimers) and detection of *MAML2*-*WT* and *GAPDH* (black arrowheads) by RT-PCR in GUMC220 and GUMC221 cells. Predicted product sizes included under each primer label. (F) Ethidium bromide-stained agarose gels showing detection of *CRTC1*-*MAML2* in H292 cell line shown as a positive control (black arrowheads). Asterisks indicate primer dimers. Predicted product sizes included under each primer label. (G) Ethidium bromide-stained agarose gels showing detection of candidate *KRT14*-*KRT5* fusion and *GAPDH* (black arrowheads) in GUMC220 and GUMC221 cells. Asterisks indicate primer dimers. PI3K, Phosphatidylinositol-4,5-bisphosphate 3-kinase. AKT, Protein kinase B. mTOR, Mechanistic Target Of Rapamycin. EGF, Epidermal Growth Factor. *EREG*, Epiregulin. *BTC*, Betacellulin. *HBEGF*, Heparin Binding EGF Like Growth Factor. *TGFA*, Transforming Growth Factor Alpha. *AREG*, Amphiregulin. *ERBB3*, Erb-B2 Receptor Tyrosine Kinase 3. *ERBB4*, Erb-B2 Receptor Tyrosine Kinase 4. *EGFR*, Epidermal Growth Factor Receptor. *ERBB2*, Erb-B2 Receptor Tyrosine Kinase 2. *CRTC1*, CREB Regulated Transcription Coactivator 1. *MAML2*, Mastermind Like Transcriptional Coactivator 2 *CRTC3*, CREB Regulated Transcription Coactivator 3. *EWS*, Ewing’s sarcoma gene. *POUFS1*, POU Class 1 Homeobox 1 M, Size marker. NTC, no template control; Neg. control, negative control (GUMC-UMB-006 (normal from tongue), Pos control, positive control.

Comparison of *EGF* and *ErbB* family member FPKM levels demonstrated that *AREG*, Heparin Binding EGF Like Growth Factor (*HBEGF*), *EGFR* and *ERBB2* were expressed at relatively higher levels than other family members in all the paired samples (Figure 3C). GUMC220/221 demonstrated differential *AREG* expression levels with statistically significantly higher levels in GUMC221 as compared to GUMC220 on both bioinformatics analysis (48,49) and quantitative Reverse Transcriptase-Polymerase Chain Reaction (qRT-PCR) (Figure 3D). Neither a bioinformatics approach (FusionCatcher, Nicorici et al. 2014) nor RT-PCR using established primer sets identified any known MEC-associated fusion gene transcripts (Figure 3E, F, Supplementary Table 1). A candidate novel fusion gene (*KRT14*-*KRT5*) was identified using FusionCatcher (Supplementary Table 2). Primers designed around the predicted fusion site revealed a product of the expected size on RT-PCR (Figure 3G, Supplementary Table 3, Supplementary Figure 2A). Sequencing of the PCR product revealed a candidate junction site (Supplementary Figure 2B) with the predicted location of the candidate fusion within exon 1 (Supplementary Figure 2C). Fluorescence *in situ* hybridization (FISH) was performed on cultured cancer cells and the original mucoepidermoid cancer to determine if K14 and K5 probes would co-localize to the same chromosome but results were not definitive (Supplementary Figure 2D, E).

### An activated AREG-EGFR-AKT pathway was confirmed as present in the original tissue and xenograft of the well-differentiated mucoepidermoid carcinoma

Presence of AREG, p-EGFR, p-AKT and p-mTOR were confirmed in the cancer tissue, CRC cells and xenografts by immunohistochemistry (IHC) (Figure 4A-C). However, although Matrigel cultures showed preservation of AREG and EGFR expression, reactivity for p-AKT and p-mTOR were lower compared to other specimens (Figure 4D). Reads Per Kilobase of transcript per Million mapped reads (RPKM) reported for normal salivary gland from the Human Protein Atlas (HPA) with FPKM levels for *AREG*, *EGFR*, *AKT* and *mTOR* from MEC GUMC220/221 are shown for comparison (Figure 4E). KRT protein expression was evaluated for concordance between original tissue and cultured cells to follow-up on the candidate *KRT14*-*KRT5* fusion gene identified in cultured cells. KRT8, KRT18 and TP53 were included for comparison because they are also expressed at relatively higher levels under CRC conditions (34). Reactivity for KRT5/18 was higher than KRT14/8 in the original MEC (Figure 5A). In contrast primary cells in CRC showed higher reactivity for KRT5/14 than KRT8/18 and this pattern was largely preserved in Matrigel culture (Figure 5B, C). Nuclear-localized TP53 expression found in the original cancer was maintained in the CRC and Matrigel cultures (Figure 5 A-C). RPKM reported for normal salivary gland from the Human Protein Atlas (HPA) with FPKM levels for *KRT5/14/8/18* and *TP53* from MEC GUMC220/221 are shown for comparison (Figure 5D).

**Figure 4.**
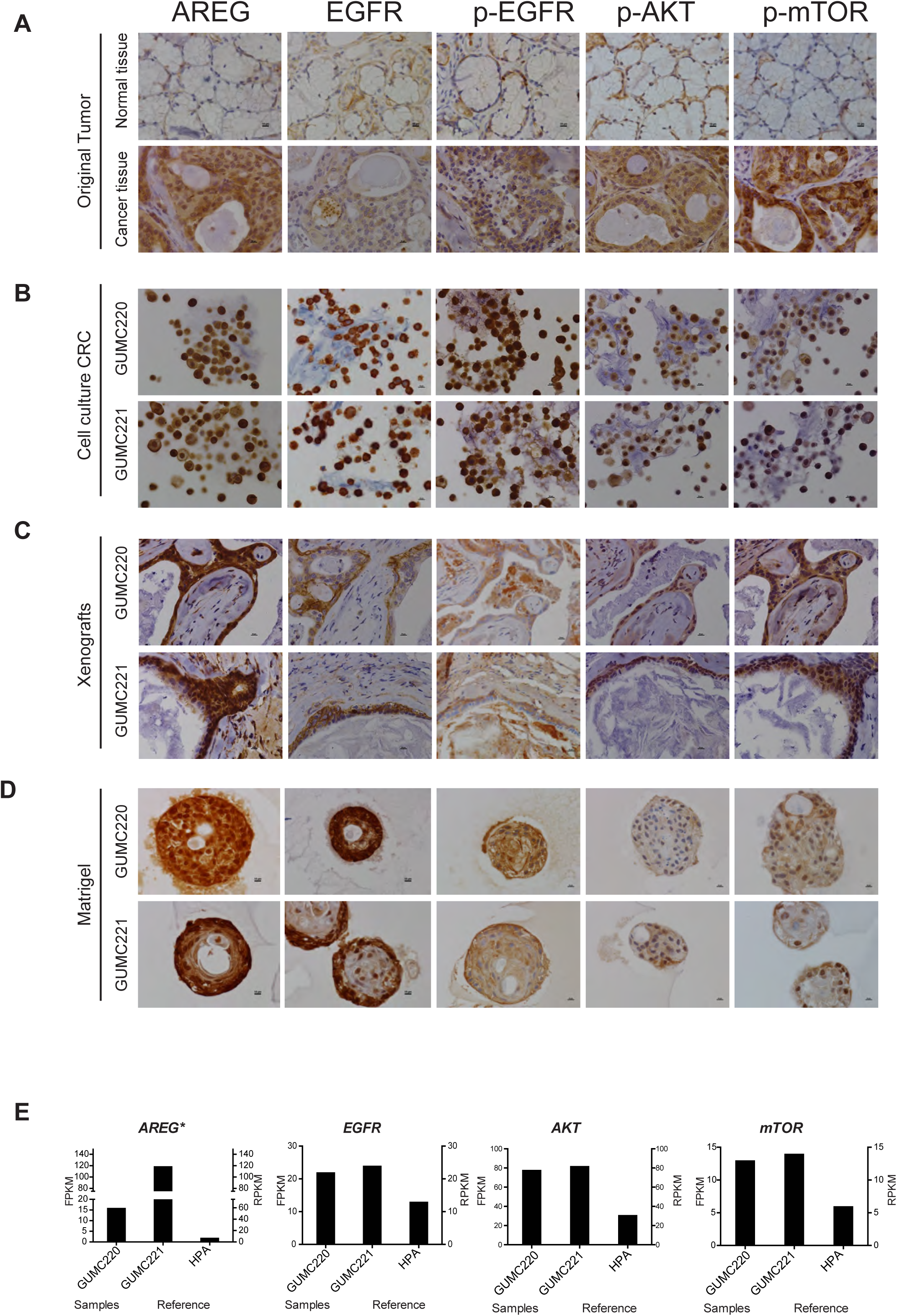
Evaluation of AREG, EGFR, p-EGFR, p-AKT, and p-mTOR expression in original tumor and GUMC220/221 CRC cultures, xenografts and Matrigel cultures. Representative IHC images for AREG, EGFR, p-EGFR, p-AKT, and p-mTOR in (A) adjacent normal tissue (top) and mucoepidermoid carcinoma (cancer tissue) (bottom), (B) GUMC220 (top) and GUMC221 (bottom) cell pellets from CRC cultures, (C) GUMC220 (top) and GUMC221 (bottom) xenografts, (D) GUMC220 (top) and GUMC221 (bottom) Matrigel spheroids. Images taken at 40x. Size bar = 10μm. (E) Left, bar graphs illustrating relative FPKM values for *AREG*, *EGFR*, *AKT*, and *mTOR* in GUMC220 and GUMC221. Right, bar graphs illustrating relative RPKM reads for *AREG*, *EGFR*, *AKT*, and *mTOR* from the Human Protein Atlas (HPA) for normal salivary gland tissue as reference. ^*^*p*<0.05 between GUMC220 and GUMC221, Cuffdiff_pval.

**Figure 5.**
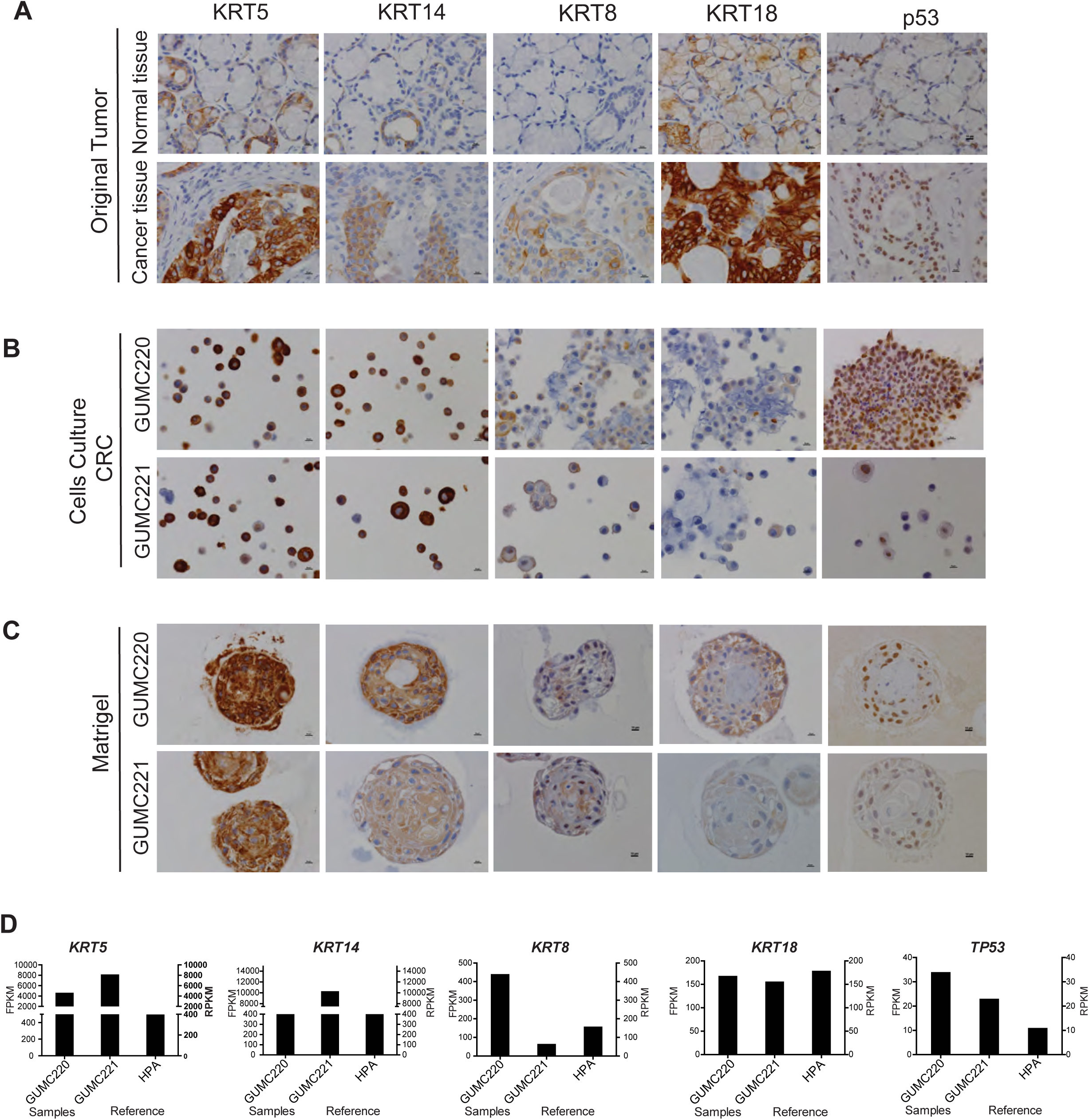
Evaluation of KRT5, KRT14, KRT8, KRT18, and p53 in original tumor and GUMC220/221 CRC and Matrigel cultures. Representative IHC images for KRT5, KRT14, KRT8, KRT18, and p53 in (A) adjacent normal tissue (top) and mucoepidermoid carcinoma (cancer tissue) (bottom), (B) GUMC220 (top) and GUMC221 (bottom) cell pellets from CRC cultures, (C) (D) GUMC220 (top) and GUMC221 (bottom) Matrigel spheroids. Images taken at 40x. Size bar = 10μm. (D) Left, bar graphs illustrating relative FPKM values for *KRT5*, *KRT14*, *KRT18*, and *Tp53* in GUMC220 and GUMC221. Right, bar graphs illustrating relative RPKM reads for *KRT5*, *KRT14*, *KRT18*, and *Tp53* from the Human Protein Atlas (HPA) for normal salivary gland tissue as reference.

### MEC GUMC220/221 cells were significantly more sensitive to AKT inhibitor MK-2206 than GSK690693

Survival of GUMC220/221 primary cells was assessed following exposure to AKT inhibitors MK2206 and GSK690693 over a range of concentrations in 2D and 3D culture conditions using the three different media shown to propagate the cells (CM+Y, CM and EpiC). Doxorubicin was used as a comparative control for the two AKT inhibitors. MDA-MD-453 cells were used as a positive control for GSK690693 (53). Survival was statistically significantly and reproducibly reduced under all conditions by MK-2206 as well as doxorubicin but not GSK690693 (Figures 6, 7). In contrast, survival of MDA-MD-453 cells was reproducibly reduced in all three media (Supplementary Figure 3A). Western blot analyses were performed to determine if the drugs induced expected changes in p-AKT, p-GSK3α/β and p-mTOR. Steady state protein levels were compared at one, two and three days of exposure to MK2206, GSK690693 and DMSO vehicle control (Figure 8). p-AKT levels were reproducibly reduced after exposure to MK2206 and increased after exposure to GSK690693 however consistent changes in p-GSK3α/β and p-mTOR were not found. In contrast, positive control MDA-MD-453 cells demonstrated the expected increase in p-AKT and reductions in p-GSK3α/β and p-MTOR following GSK690693 exposure (Supplementary Figure 3B). Because ERBB2 (HER2) over-expression has been linked to the differential pattern of MK2206 and GSK690693 sensitivity found here (29), RNA and protein expression levels were evaluated in GUMC220/221 but no increases were found (Figure 3C, Supplementary Figure 4). Because reduced *AKT3* expression levels in triple negative breast cancers are reported to increase sensitivity to GSK690693 (Chin et al. 2014), FPKM values of *AKT1/2/3* were checked in GUMC 220/221. FPKM levels were highest for *AKT1* (82.28/77.81) followed by *AKT2* (27.25/22.38) and *AKT3* (1.17/1.62).

**Figure 6.**
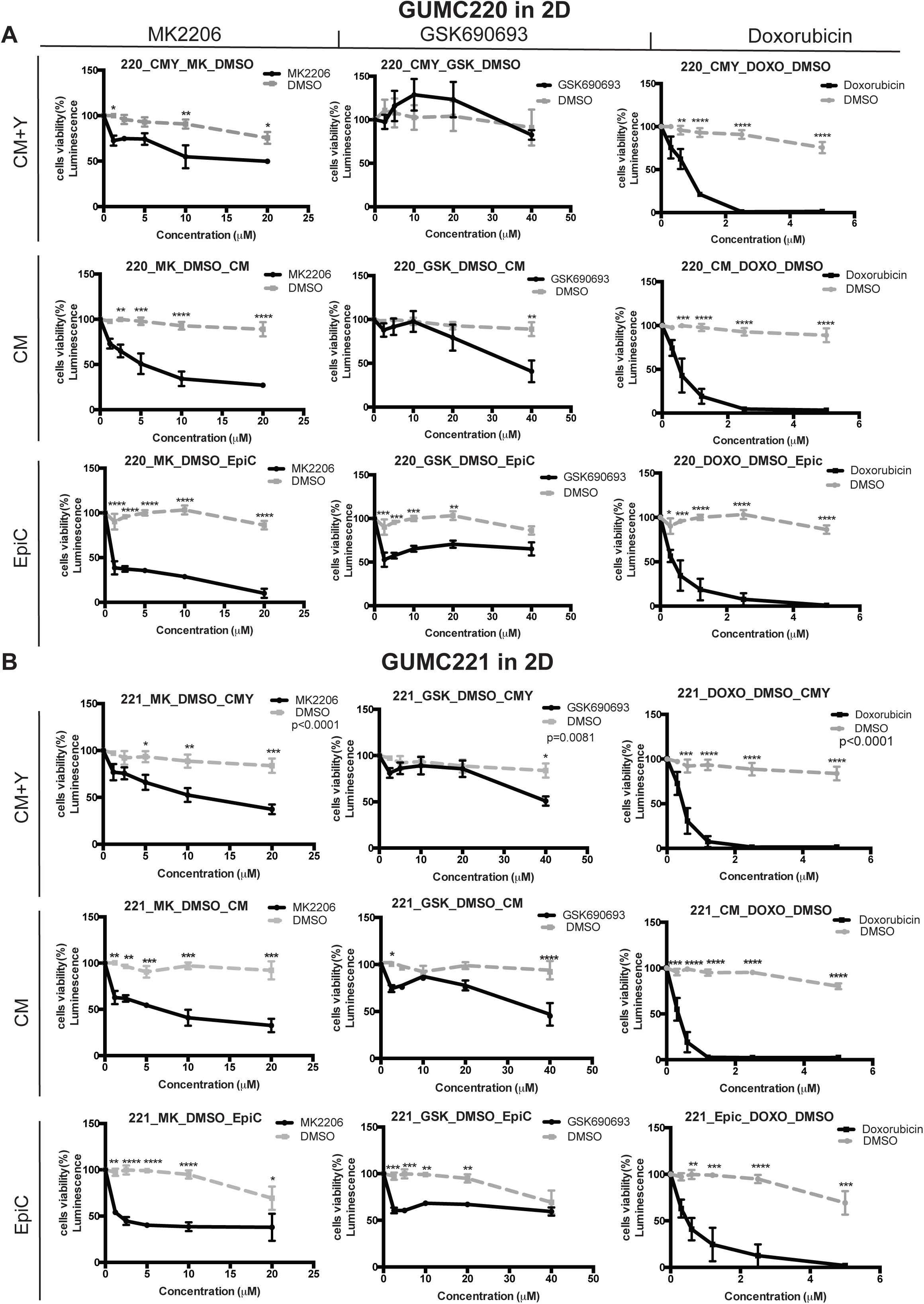
Viability of GUMC220/221 2D cell cultures exposed to MK2206, GSK690639 and doxorubicin in CM+Y, CM and EpiC media. Cell viability measured after three days for GUMC220 (A) and GUMC221 (B) under three different 2D culture media (CM+Y, CM, and EpiC) at a range of concentrations in the presence of left, MK2206 (1.2μM-20μM, black line) compared to DMSO (vehicle control, gray line), middle, GSK690639 (2.5μM-40μM, black line) compared to DMSO (vehicle control, gray line), right, doxorubicin (0.3μM-5μM, black line) compared to DMSO (vehicle control, gray line). Mean ± SEM shown for each concentration. Experimental design used one DMSO vehicle control for each media (dotted grey line) each time experiment performed, n=3. ^*^ *p*≤ *0.05*, ^**^*p*≤ 0.01, ^***^*p*≤0.001, ^****^*p*≤0.0001, One-way ANOVA.

**Figure 7.**
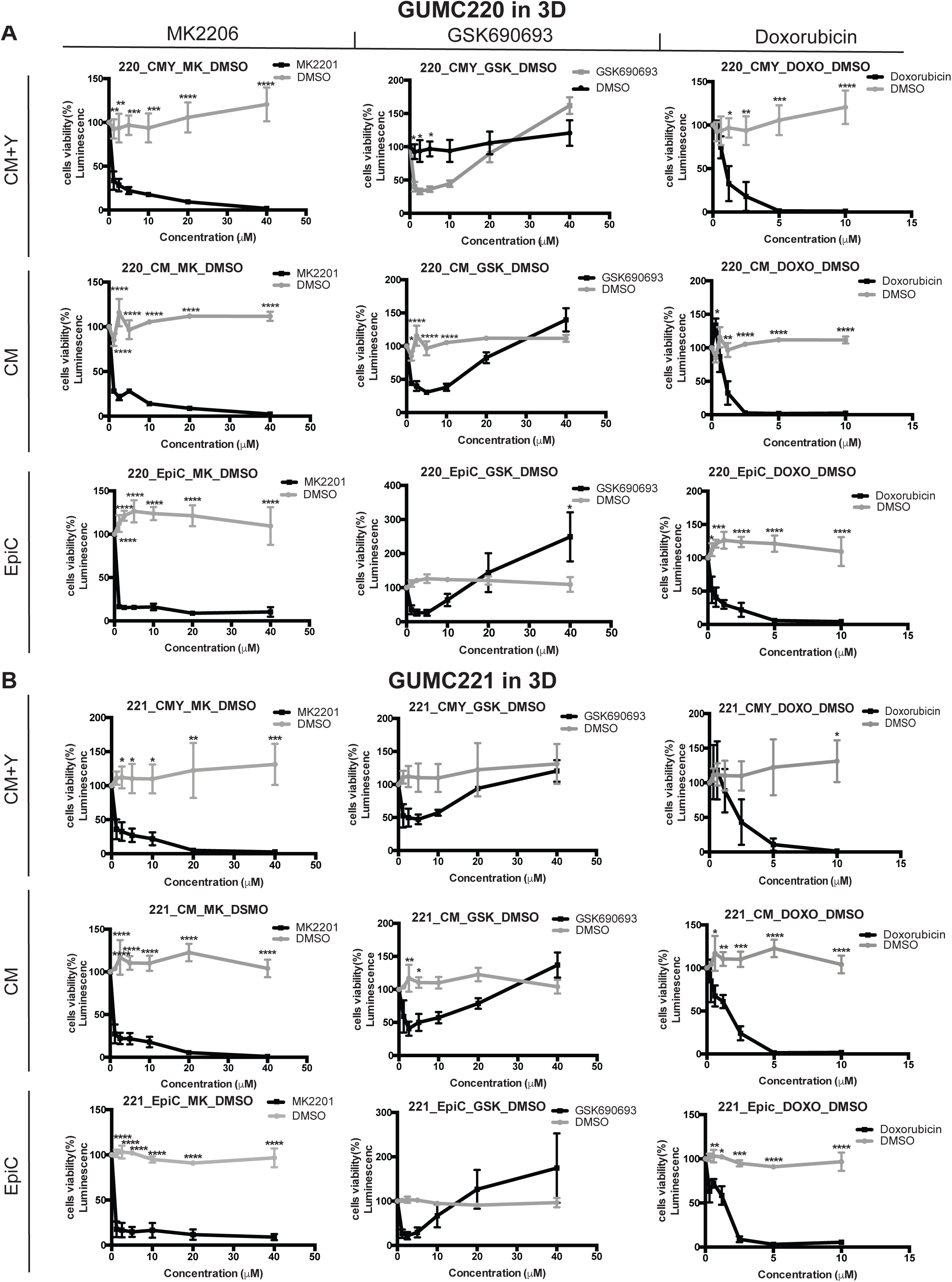
Viability of GUMC220/221 3D cell cultures exposed to MK2206, GSK690639 and doxorubicin in CM+Y, CM and EpiC media. Cell viability measured after three days for GUMC220 (A) and GUMC221 (B) under three different 3D culture media (CM+Y, CM, and EpiC) at a range of concentrations in the presence of left, MK2206 (1.2μM-40μM, black line) compared to DMSO (vehicle control, gray line), middle, GSK690639 (2.5μM-40μM, black line) compared to DMSO (vehicle control, gray line), right, doxorubicin (0.6μM-10μM, black line) compared to DMSO (vehicle control, gray line). Mean ± SEM shown for each concentration. Experimental design used one DMSO vehicle control for each media (dotted grey line) each time experiment performed, n=3. ^*^ *p*≤ *0.05*, ^**^*p*≤ 0.01, ^***^*p*≤0.001, ^****^*p*≤0.0001, One-way ANOVA.

**Figure 8.**
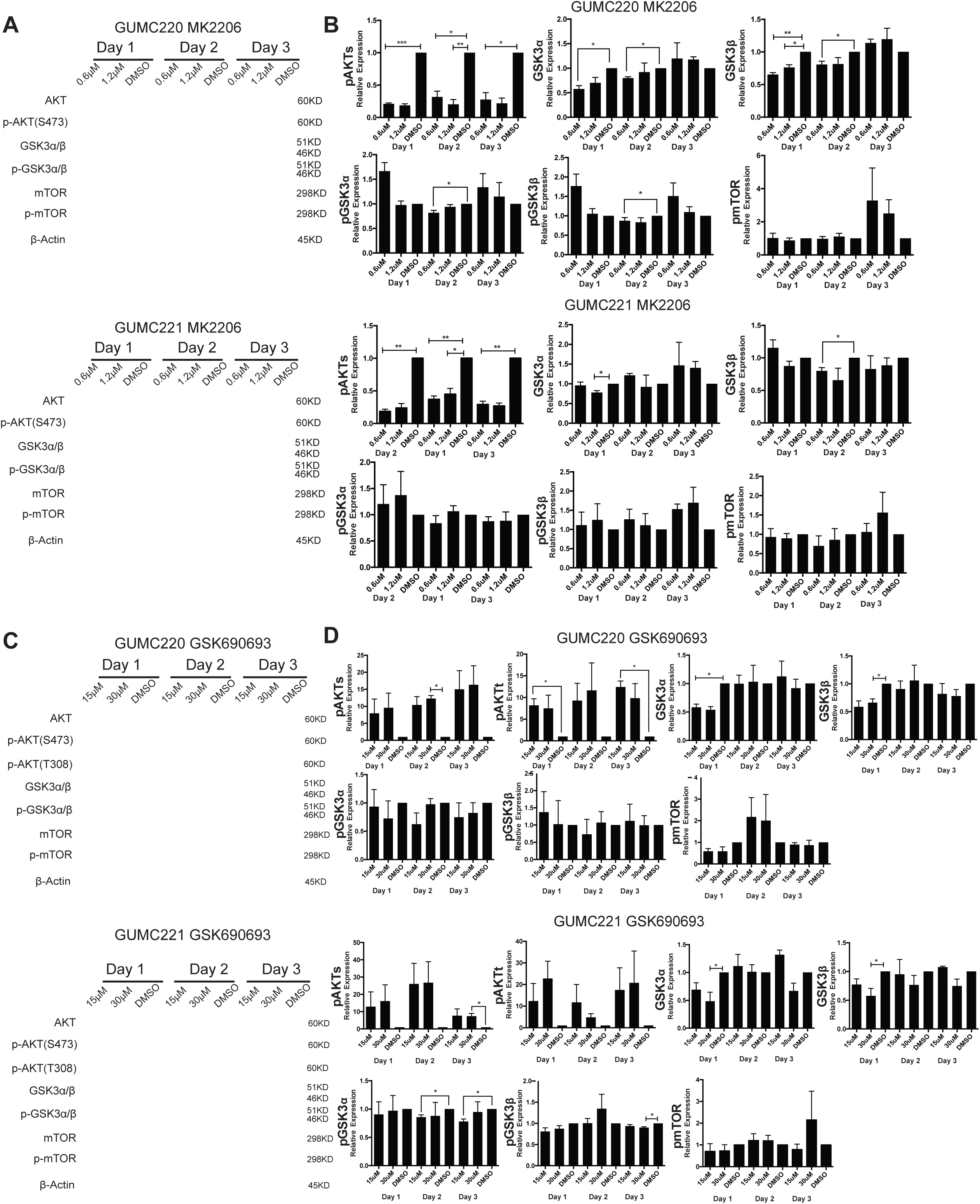
Western blot analyses comparing three day time-course of steady state levels of AKT, pAKT, GSK3α/β, p-GSK3α/β, mTOR, and p-mTOR following exposure to MK2206 and GSK690693 in 2D culture (CM+Y). (A) Representative western blots of AKT, p-AKT(S473), GSK3α/β, p-GSK3α/β, mTOR, p-mTOR and β-Actin from GUMC220 and GUMC221 exposed to MK2206 (0.6μM and 1.2μM) compared with DMSO vehicle control. (B) Bar graphs illustrating the timecourse of changes in steady state expression levels of pAKT(S473), GSK3α/β, p-GSK3α/β, and p-mTOR in GUMC220 and GUMC221 cells after 1,2 and 3 days exposure to the two different concentrations of MK2206 and DMSO. (C) Representative western blots of AKT, p-AKT(S473), p-AKT(T308), GSK3α/β, p-GSK3α/β, mTOR, p-mTOR, and β-Actin from GUMC220 and GUMC221 under 2D CM+Y culture exposed to GSK690693 (15μM and 30μM) compared with DMSO vehicle control. (D) Bar graphs illustrating the timecourse of changes in steady state expression levels of pAKT(S473), p-AKT(T308), GSK3α/β, p-GSK3α/β, and p-mTOR in GUMC220 and GUMC221 cells after 1, 2 and 3 days exposure to the two different concentrations of GSK690693 compared with DMSO vehicle control. Phosphoprotein levels were normalized to total proteins that were normalized to actin. ^*^ *p*≤ 0.05, ^**^*p*≤ 0.01, ^***^*p*≤0.001, unpaired t-test one tailed. Protein lysates collected after 1, 2 and 3 days of drug exposure. Mean ± SEM shown for each concentration and DMSO, n=3. Supplemental figures.

## Discussion

Primary cell cultures were established from all of the benign, primary cancer and metastatic samples attempted although not the two chronic sialoadenitis cases. Chronic sialoadenitis is an inflammatory disease that leads to atrophy and loss of acinar structures (54). It is possible that either or both the presence of inflammation or diminished epithelia contributed to the lack of success with this disease. Epithelial cells were isolated from a salivary gland containing a lymphoma (GUMC311/312) demonstrates an approach for isolating nonmalignant epithelial cells from the lymphoma environment for studies targeted towards the study of heterogeneous cell populations in malignancy. MEC GUMC220/221 cultures were able to form a patient derived xenograft but this was not true of all cultures attempted. The same has been reported for CRC-derived cells from human pancreatic cancers (55). It could be notable that only the MEC GUMC220/221 cells carried a cancer-risk associated *TP53* SNP altering p53 function (56). CRC-derived murine cancer cells with *Trp53* haploinsufficiency effectively form allografts (34).

The primary human cultures established using the classic CRC technology were secondarily propagated in alternative media, as reported previously for murine cells (34). They also propagated in 3D culture formats. 3D culture formats have been proposed as being more representative of the *in vivo* setting leading to more accurate predictive chemosensitivity testing (57). Here the differential sensitivity of the two AKT inhibitors was present in both 2D and 3D culture formats. It also was present across the different culture media tested. Because rho kinases are linked to multiple roles in carcinogenesis (58) and Rho kinase inhibitors impact cellular physiology in several ways (59,60), a concern was whether or not the presence of Y-27632 would distort chemotherapeutic testing. But this was not the case for the agents tested, MK2206, GSK690693, doxorubicin, as was previously shown for Vorinostat tested in CRC cells from an HPV11-related laryngeal carcinoma (32).

The differential response to the two different AKT inhibitors underscores the more general need for a further understanding of the genetic settings in which specific anti-cancer drugs will be active. The relative resistance to GSK690693 in the MEC cells appeared to be innate rather than acquired in culture as it was evident on first testing and found in all culture formats. Significantly previously described mechanisms of GSK690693 resistance were not present. In the clinic AKT inhibitors are anticipated to be used in combination with other chemotherapeutic agents, not as single agents (24). One next step would be to evaluate response to MK2206 in rational combinations with other agents active in salivary gland cancers (61). More generally, the approach described here can be used to expand testing of human primary cancer cells to further our knowledge of chemosensitivity of salivary gland cancers.

Reproducible results from RNAseq and exome sequencing were obtained from cells established in CRC at low passage, providing data specifically on the epithelial component of the tissue and mitigating variables associated with more prolonged cell culture including gene expression drift (34). There was high concordance on genetic findings from each of the paired samples, even though they were derived from were geographically distinct sites within the tumors. This could argue that single specimens may be sufficient. At the same time the high levels of AREG expression found in the MEC reported here were only present in one of the two primary cultures, perhaps arguing against single specimens for precision medicine.

In salivary gland cancers fusion genes are seen as key molecular drivers (16). In MEC they are reported to be more frequently present in low-grade MEC and in association with AREG overexpression but that was not the case for the MEC studied here. The presence of a novel *KRT14*-*KRT5* fusion gene here could not be fully confirmed at the DNA level. FPKM values for KRT14 and KRT5 were relatively high in the cultured cells. It is possible the fusion detected could be a chimeric RNA or a spurious finding as has been previously reported for RNAseq data (62,63).

In summary, the approach employed here can reasonably, rapidly and cost-effectively expand primary cell cultures from salivary gland neoplasms for genetic and chemosensitivity studies, extending our understanding of the pathophysiology of these relatively uncommon cancer types and providing a foundation for precision medicine.

## Acknowledgments

All authors have read the journal’s policy on disclosure of potential conflicts of interest. All authors have disclosed any financial or personal relationship with organizations that could potentially be perceived as influencing the described research. AMA, JKB, BRH, WW, XZ, SC, EK, BKA BJD, & PAF declare no conflict of interest. XL declares that Georgetown University has submitted a patent application for the cell reprogramming technology described, on which X Liu is an inventor. The intellectual property is under an exclusive license to Propagenix, in which Georgetown University has a founding equity interest. We thank GUMC core facilities NTSR for patient recruitment, GESR for preparation of DNA and RNA for sequencing, and HTSR for performing IHC. Research supported by NIH NIDCR 1R56DE023259 (PAF), NIH NCI 5P30CA051008 (Histopathology and Tissue, Genomics and Epigenomics, Tissue Culture, and Animal Shared Resources), and King Khalid University, Abha, Saudi Arabia (AMA). Authorship contributions: conception and design (AMA, XL, BJD, PAF), data acquisition (AMA, JKB, BRH, WW, SC, EK, PAF), analysis and interpretation of data (AMA, JKB, BRH, XZ, BVK, PAF), manuscript preparation (AMA, XL, JKB, BRH, WW, XZ, SC, EK, BVK, BJD, PAF).

---

**Supplementary Figure 1**

**Dendrogram illustrating hierarchical clustering of paired culture transcriptomes.** Mucoepidermoid carcinoma (GUMC220/221). Metastatic poorly differentiated squamous cell carcinoma (GUMC264/265). Benign pleomorphic adenoma (GUMC572/573, GUMC332/349, GUMC436/446, GUMC284/299). Salivary gland containing lymphoma (GUMC311/312).

**Supplementary Figure 2**

***KRT14*-*KRT5* gene fusion analyses.** (A) Ethidium bromide stained agarose gel illustrating RT-PCR screening for eight primer sets, *KRT5*-*KRT13*, *KRT5*-*KRT14*, *KRT5*-*KRT14*_*1*, *KRT5*-*KRT14*_*2*, *KRT5*-*NCL*, *KRT14*-*KRT5*_*1*, *KRT14*-*KRT5*_*2*, *KRT14*-*KRT5*_*3*, targeting the potential fusion gene junction identified by FusionCatcher in GUMC220 and GUMC221. Arrowhead indicates PCR products isolated for sequencing from the *KRT14*-*KRT5*_*1* primer pair. (B) Base-calling sequencing electropherogram of the *KRT14*-*KRT5*_*1* RT-PCR amplicon illustrating potential *KRT14*-*KRT5* fusion gene junction sequence in GUMC220 and GUMC221. (C) Predicted exon 1 site of *KRT14*-*KRT5* fusion gene junction within predicted exon structure of fusion gene shown with *KRT14* gene exon structure (left) and *KRT5* gene exon structure (right). (D) Fluorescence images illustrating *KRT14* (red) and *KRT5* (green) FISH performed on chromosomes harvested from CRC-cultured GUMC220 and GUMC221 cells. Images taken at 100X. Top left insert shows magnified images. (E) Fluorescence images illustrating *KRT14* (red) and *KRT5* (green). FISH performed on formalin fixed paraffin embedded (FFPE) section from the mucoepidermoid carcinoma GUMC220/221 were derived from. Images taken at 100X. Top left insert shows magnified images. Yellow arrows indicted probes.

**Supplementary Figure 3**

**Viability of MD-MB-453 2D cell cultures exposed to MK2206, GSK690639 and doxorubicin in CM+Y, CM and EpiC media with western blot analyses of steady state levels of AKT, p-AKT, GSK3α/β, p-GSK3α/β, mTOR, and p-mTOR following exposure to GSK690693 in CM+Y.** (A) Cell viability measured after three days for MD-MB-453 cells under three different 2D culture media (CM+Y, CM, and EpiC) at a range of concentrations in the presence of left, MK2206 (1.2μM-20μM, black line) compared to DMSO (vehicle control, gray line), middle, GSK690639 (2.5μM-40μM, black line) compared to DMSO (vehicle control, gray line), right, doxorubicin (0.3μM-5μM, black line) compared to DMSO (vehicle control, gray line). Mean ± SEM shown for each concentration. Experimental design used one DMSO vehicle control for each media (dotted grey line) each time experiment performed, n=3. ^*^ *p*≤ 0.05, ^**^*p*≤ 0.01, ^***^*p*≤0.001, ^****^*p*≤0.0001, One-way ANOVA. (B) Representative western blots of steady state levels of AKT, p-AKT(S473), p-AKT(T308), GSK3α/β, p-GSK3α/β, mTOR, p-mTOR, and β-Actin from MDA-MB-453 cells under 2D CM+Y culture exposed to two different concentrations of GSK690693 (15μM and 30μM) compared with DMSO-only control. Protein lysates collected after 3 days of drug exposure. KD, Kilodaltons.

**Supplementary Figure 4**

**Immunohistochemistry for HER2.** (A) Representative HER2 IHC image of human breast cancer tissue used as a positive control. (B) Representative HER2 IHC images of adjacent normal tissue (top) and mucoepidermoid carcinoma cancer tissue (bottom). (C) Representative HER2 IHC images of GUMC220 (top) and GUMC221 (bottom) cell pellets from CRC cultures.

